# Sterilizing Immunity against SARS-CoV-2 Infection in Mice by a Single-Shot and Modified Imidazoquinoline TLR7/8 Agonist-Adjuvanted Recombinant Spike Protein Vaccine

**DOI:** 10.1101/2020.10.23.344085

**Authors:** Sonia Jangra, Jana De Vrieze, Angela Choi, Raveen Rathnasinghe, Gabriel Laghlali, Annemiek Uvyn, Simon Van Herck, Lutz Nuhn, Kim Deswarte, Zifu Zhong, Niek Sanders, Stefan Lienenklaus, Sunil David, Shirin Strohmeier, Fatima Amanat, Florian Krammer, Hamida Hammad, Bart N. Lambrecht, Lynda Coughlan, Adolfo García-Sastre, Bruno G. De Geest, Michael Schotsaert

**Author notes:** Contributed equally (Sonia Jangra, Jana De Vrieze, Angela Choi). University of Maryland School of Medicine, Department of Microbiology and Immunology and Center for Vaccine Development and Global Health (CVD), Baltimore, MD,USA.

## Abstract

The search for vaccines that protect from severe morbidity and mortality as a result of infection with severe acute respiratory syndrome coronavirus 2 (SARS-CoV-2), the virus that causes coronavirus disease 2019 (COVID-19) is a race against the clock and the virus. Several vaccine candidates are currently being tested in the clinic. Inactivated virus and recombinant protein vaccines can be safe options but may require adjuvants to induce robust immune responses efficiently. In this work we describe the use of a novel amphiphilic imidazoquinoline (IMDQ-PEG-CHOL) TLR7/8 adjuvant, consisting of an imidazoquinoline conjugated to the chain end of a cholesterol-poly(ethylene glycol) macromolecular amphiphile). This amphiphile is water soluble and exhibits massive translocation to lymph nodes upon local administration, likely through binding to albumin. IMDQ-PEG-CHOL is used to induce a protective immune response against SARS-CoV-2 after single vaccination with trimeric recombinant SARS-CoV-2 spike protein in the BALB/c mouse model. Inclusion of amphiphilic IMDQ-PEG-CHOL in the SARS-CoV-2 spike vaccine formulation resulted in enhanced immune cell recruitment and activation in the draining lymph node. IMDQ-PEG-CHOL has a better safety profile compared to native soluble IMDQ as the former induces a more localized immune response upon local injection, preventing systemic inflammation. Moreover, IMDQ-PEG-CHOL adjuvanted vaccine induced enhanced ELISA and *in vitro* microneutralization titers, and a more balanced IgG2a/IgG1 response. To correlate vaccine responses with control of virus replication *in vivo*, vaccinated mice were challenged with SARS-CoV-2 virus after being sensitized by intranasal adenovirus-mediated expression of the human angiotensin converting enzyme 2 (ACE2) gene. Animals vaccinated with trimeric recombinant spike protein vaccine without adjuvant had lung virus titers comparable to non-vaccinated control mice, whereas animals vaccinated with IMDQ-PEG-CHOL-adjuvanted vaccine controlled viral replication and infectious viruses could not be recovered from their lungs at day 4 post infection. In order to test whether IMDQ-PEG-CHOL could also be used to adjuvant vaccines currently licensed for use in humans, proof of concept was also provided by using the same IMDQ-PEG-CHOL to adjuvant human quadrivalent inactivated influenza virus split vaccine, which resulted in enhanced hemagglutination inhibition titers and a more balanced IgG2a/IgG1 antibody response. Enhanced influenza vaccine responses correlated with better virus control when mice were given a lethal influenza virus challenge. Our results underscore the potential use of IMDQ-PEG-CHOL as an adjuvant to achieve protection after single immunization with recombinant protein and inactivated vaccines against respiratory viruses, such as SARS-CoV-2 and influenza viruses.

## Introduction

Severe acute respiratory syndrome coronavirus 2 (SARS-CoV-2), as the causative agent of coronavirus disease 2019, also known as COVID-19, is a beta-coronavirus which belongs to the family of *Coronaviridae* and is currently responsible for the third human coronavirus outbreak in the past 20 years, after SARS (now often referred to as SARS-CoV-1) in 2002/03 and MERS (Middle east respiratory syndrome) in 2012 (1). SARS-CoV-2 was first identified in Wuhan, China in December 2019 (2,3). This COVID-19 pandemic has caused unprecedented morbidity, mortality and global economic instability. SARS-CoV-2 is highly pathogenic and is believed to spread mainly through respiratory droplets and aerosols. The current preventive measures include quarantine, isolation and physical social distancing. Thus far, therapeutic drugs are of limited use in the clinic, and no specific vaccine is available yet, therefore calling for an urgent need for development of effective vaccines to restrict disease as well as viral spread.

More than hundred candidate vaccines, consisting of multiple vaccine types such as recombinant viral epitopes (surface glycoprotein), adenovirus-based vectors (e.g. recombinant replication incompetent HAdV-C5), purified inactivated or live-attenuated virus, virus like particles (VLPs) and DNA or RNA based vaccine formulations, are currently being investigated (4,5). At present mRNA-based vaccines formulated in lipid nanoparticles, recombinant protein-based and inactivated virus-based vaccines as well as viral vector-based vaccines have reached late stage of clinical development, entering phase 3 testing. For these vaccines, pre-clinical data in animal models has also been generated supporting the hypothesis that these vaccines can effectively prevent viral infection. However, little is known about whether recombinant protein vaccines are capable of conferring protective immunity. In contrast to the aforementioned mRNA and viral vector-based vaccines, recombinant protein vaccines are simpler as they consist of a single entity antigen and – in contrast to viral vectors – do not require antigen expression in the vaccinees. Compared to mRNA vaccines, recombinant protein vaccines do not require complex (lipid) nanoparticle formulations to overcome the formidable barrier of the endosomal membrane before reaching the cytoplasm which is the subcellular target compartment for the antigen-expressing mRNA. Moreover, thus far no mRNA-based vaccine has been licensed, which might pose additional hurdles in view of mass manufacturing in world-wide immunization campaigns.

Hence, exploring the viability of a recombinant protein COVID-19 vaccine might be of considerable relevance. SARS-CoV-2 consists of over 30 kb single-stranded positive strand RNA genome which encodes four major structural proteins, spike (S), membrane (M), nucleocapsid (N) and envelope (E). The spike protein comprises a homotrimeric structure which is present on the surface of the virus and facilitates the viral attachment and entry into the host cells. Like SARS-CoV-1, SARS-CoV-2 S protein gains entry into host cells via human angiotensin-converting enzyme 2 (hACE-2) receptors on the host cell surface via its receptor-binding domain (RBD) (1)(6). Subsequently, membrane-associated serine proteases such as transmembrane protease, serine 2 (TMPRSS2) or endosomal-associated proteases such as cathepsins cleave the S protein, thereby promoting efficient fusion of the viral membrane to the host cell membrane, followed by release of viral content into the cell cytoplasm, where the virus subsequently replicates. The viral infection usually begins in the oral/nasal cavity and once released, it gradually establishes itself in type-II pneumocytes of the lower respiratory air tract and enterocytes in the gastrointestinal tract (7,8).

Due to its involvement in viral entry, the S protein is a major target for current vaccine development against SARS-CoV-2 (5). Therefore, in this study we explored the recombinant SARS-CoV-2 S protein as a potential vaccine candidate. As recombinant protein antigens are poorly immunogenic and are incapable of mounting antigen-specific immunity of sufficient quality, amplitude and duration, co-administration of adjuvants that shape B cell and T cell responses are indispensable. Adjuvants like alum and oil-in-water emulsions can act through a multitude of mechanisms. More defined small molecule adjuvants that potently activate innate immune cells by triggering specific innate immune receptors might be more relevant for anti-viral vaccine design. The Toll-like receptors 7 and 8 (TLR7/8) are widely distributed amongst innate immune cell subsets over a broad range of species (9). Akin to be an endosomal pattern recognition receptor for viral RNA, triggering of these receptors provokes robust type I interferon production that can skew a Th1-type adaptive immune response against co-administered antigen (10). The latter are characterized by robust antibody titers capable of inducing viral neutralization through a variety of mechanisms, including Fc-mediated innate immune killing as well as inducing CD4- and CD8 T-cell based immunological memory. Moreover, vaccines adjuvanted with TLR7/8 ligands have been shown to confer enhanced protective immunity in both mouse and non-human primate models (11,12).

Being well-defined small molecules, imidazoquinolines are a class of TLR7/8 agonists (13) that hold a massive technological advantage in terms of production and physicochemical stability. However, their pharmacokinetic profile is characterized by rapid systemic dissemination upon local (e.g. subcutaneous or intramuscular) administration, thereby causing unwanted innate immune activation at multiple distal tissues (14), which is currently a strong limitation for applying imidazoquinoline TLR7/8 agonists in mass immunization campaigns. We and others have reported on strategies to alter the bio-distribution of imidazoquinolines through chemical conjugation to a synthetic carrier that limits systemic circulation but confers robust translocation to immune-inducing sites in sentinel lymph nodes (14–18). In the present work, we report on a novel amphiphilic carrier for imidazoquinoline (IMDQ) TLR7/8 agonists with high translational potential, based on conjugation of a single imidazoquinoline to the chain end of a cholesteryl-polyethylene glycol macromolecular amphiphile (IMDQ-PEG-CHOL; Figure 1A). This design mediates binding to serum proteins such as albumin (15,19) (Figure 1B) and in contrast to pure lipidation, the conjugate is well water-soluble. As a result, mobility in tissue is achieved without depot formation, with potential loss in adjuvant efficacy, hence avoiding the need for additional formulation.

**Figure 1.**
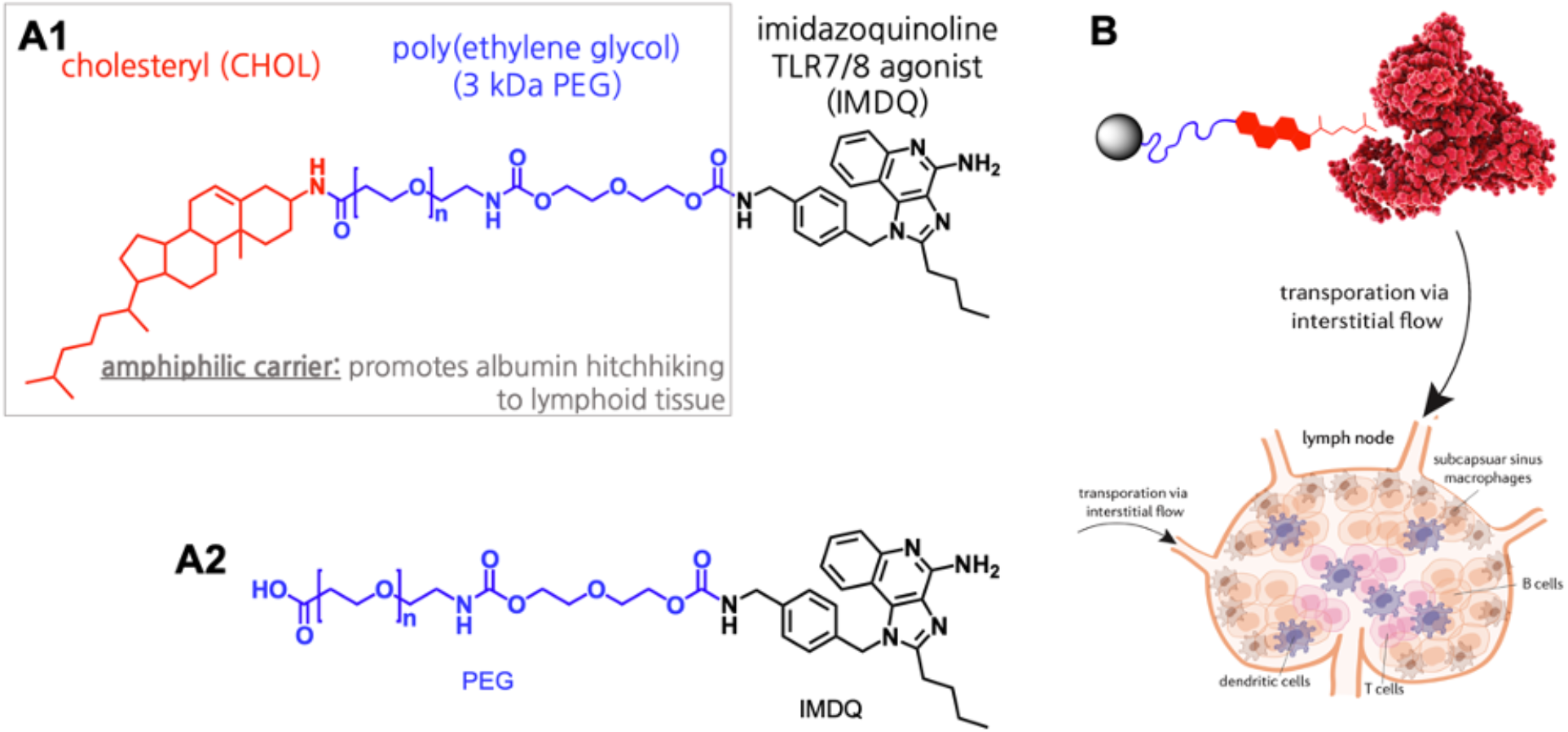
**(A)** Molecular structure of (A1) CHOL-PEG-IMDQ and (A2) PEG-IMDQ. Conjugation was performed by amide bond formation between respectively cholesterylamine and PEG and PEG and IMDQ. **(B)** Schematic representation of albumin hitchhiking-mediated lymphatic transportation.

Here we report that IMDQ-PEG-CHOL is a potent adjuvant candidate which can enhance vaccine efficiency and induce robust Th1 skewed antibody responses in mice when delivered as a single shot with either admixed S protein (for SARS-CoV-2) or seasonal quadrivalent inactivated influenza virus vaccine (QIV, for influenza). Moreover, IMDQ-PEG-CHOL was able to infer protection in SARS-CoV-2 or influenza virus (H1N1)-infected infected mice. In this context, it is noteworthy to mention that a major limitation in COVID19-vaccine development is the lack of susceptible small animal models for pre-clinical assessment and evaluation of its efficacy. Reportedly, in contrast to human ACE-2, murine ACE-2 is not targeted by wild type SARS-CoV-2 virus because of species-specific variations in ACE-2 receptors between mouse and human. Here we made use of a mouse model where hACE-2 was introduced through an adenoviral vector (Ad5-hACE2), which allows for subsequent replication of SARS-CoV-2 upon infection in the airways of transduced mice (20).

## Results and Discussion

### IMDQ-PEG-CHOL induces potent innate immune activation in lymph nodes

The imidazoquinoline 1-(4-(aminomethyl)benzyl)-2-butyl-1H-imidazo[4,5-c]quinolin-4-amine (IMDQ) (21) was conjugated to cholesteryl-poly(ethylene glycol) (CHOL-PEG), yielding IMDQ-PEG-CHOL. As a control, non-amphiphilic IMDQ-PEG was synthesized. (Figure 1A) PEG with a molecular weight of 3 kDa was chosen as an optimal compromise between water solubility and drug load. Characterization of the conjugate was performed by matrix assisted laser desorption/ionization – time of flight (MALDI-ToF) (Supplementary Figure S1) analysis whereas high pressure liquid chromatography (HPLC) analysis (Supplementary Figure S2) proved absence of free soluble non-conjugated IMDQ. Both IMDQ-PEG-CHOL and IMDQ-PEG were watersoluble, but only IMDQ-PEG-CHOL showed affinity towards albumin as measured by biolayer interferometry (Figure 2A). On the *in vitro* level, the presence of the CHOL motif dramatically improved cellular uptake by DC2.4 (Figure 2B-C; note that for imaging purpose, IMDQ was replaced by the fluorescent probe Cynanine5), a murine model mouse dendritic cell line, and was more potent in inducing NF-κB activation in a reporter cell line (Figure 2D), while being nontoxic within the tested experimental window (Figure 2E). We attribute this to the ability of the cholesterol motif to interact with the phospholipid cell membrane.

**Figure 2.**
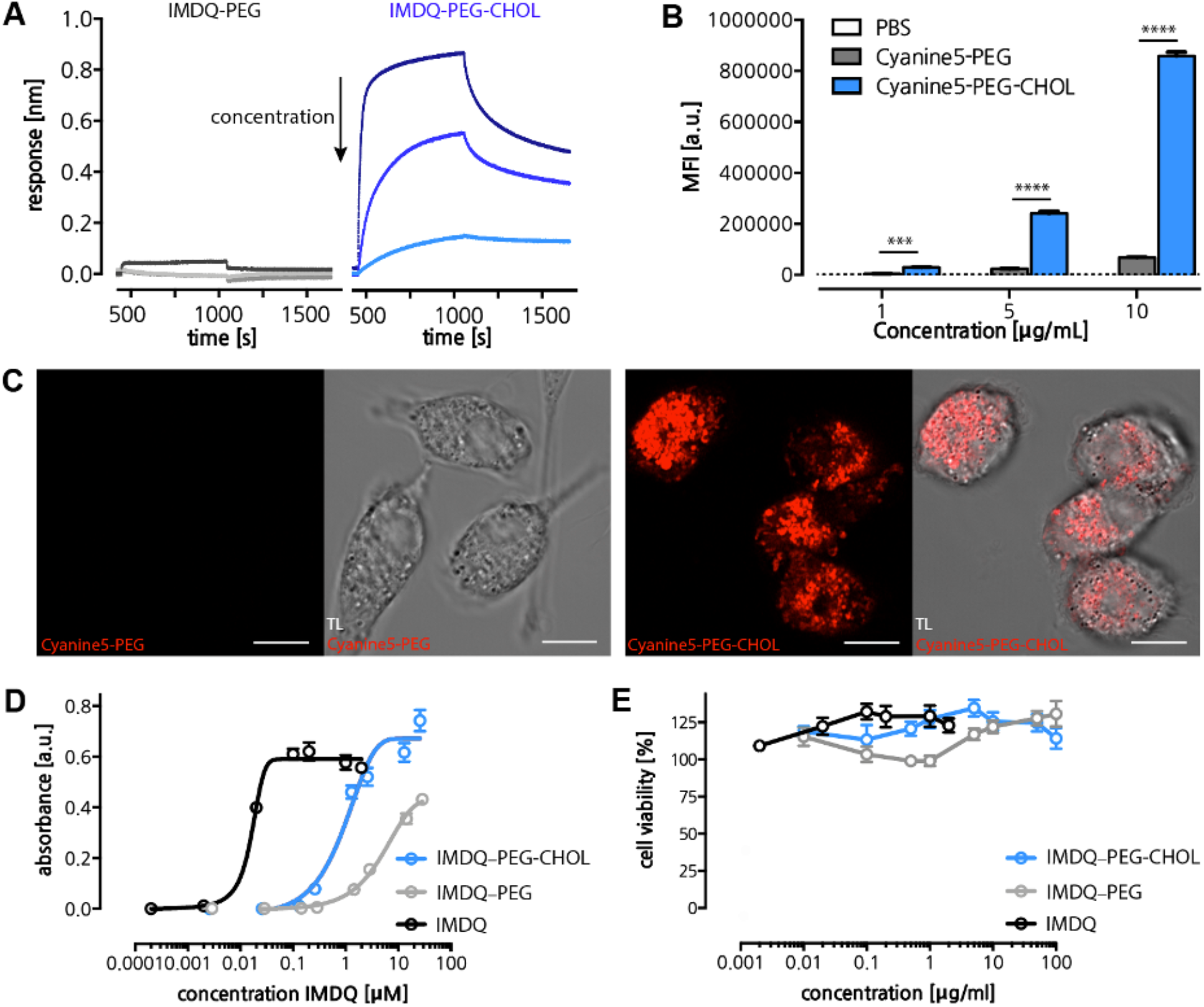
**(A)** Biolayer interferometry (BLI) sensorgrams of non-amphiphilic IMDQ-PEG and amphiphilic IMDQ-CHOL-PEG binding to albumin-coated sensors. A dilution series of 50, 10 and 5 mg/mL (dark to light color code, as marked by the black arrow) was measured. Sensors were dipped into a PEG-or lipid-PEG solution at the 500 s time point, which marks the onset of adsorption. At the 1125 s time point, sensors were dipped into PBS, which marks the onset of desorption. **(B)** Flow cytometry analysis of association between DC2.4 cells and Cyanine5-PEG, Cyanine5-PEG-CHOL, respectively. (n=3; Student t-test, ****:p<0.0001, ***:p<0.001) **(C)** Confocal microscopy images of DC2.4 cells incubated for 24 h at 37 C with Cyanine5-PEG and Cyanine5-PEG-CHOL. Left panel represents the Cyanine5 channel, right panel represents the overlay of the Cyanine5 and transmitted light channels. Scale bar represents 15 micron. **(D)** TLR agonistic activity of IMDQ-PEG-CHOL, IMDQ-PEG and native IMDQ measured as NF-κB activation using the RAWBlue reporter cell assay. (n = 6, mean + SD). **(E)** Cytotoxicity of IMDQ-PEG-CHOL, IMDQ-IMDQ and native IMDQ, measured by MTT assay (n = 6, mean + SD).

On the *in vivo* level, using a transgenic luciferase-reporter mouse model for IFN β-production (22), we found that local administration (i.e. subcutaneous injection into the footpad) of IMDQ-PEG-CHOL, in contrast to unformulated IMDQ, dramatically reduced systemic innate immune activation, while focusing its activity to the site of injection and the draining (popliteal) lymph node (Figure 3A). For quantification of the luminescence imaging data we refer to Supplementary Figure S3. Interestingly, the CHOL motif appeared crucial for mediating lymphatic translocation as the IMDQ-PEG control induced very limited activity in the draining lymph node. To further support this, we performed microscopic (Figure 3B1) and flow cytometry (Figure 3B2) analysis of popliteal lymph nodes of mice that received fluorescent Cyanine5-PEG-CHOL or Cyanine-PEG, respectively. These experiments revealed a dramatic increase in fluorescence when the conjugates contained the CHOL motif. A more detailed analysis of immune cells subsets in the draining lymph node revealed that vast percentages of lymphocytes, notably over 50% of dendritic cells (DCs) and macrophages (Mf), as well as 40% of B cells, were targeted by Cyanine5-PEG-CHOL (Figure 3C). In a similar experimental setting, IMDQ-PEG-CHOL induced recruitment (Figure 3D1) and robust activation (Figure 3D2) of immune cells. Taken together, these data support our hypothesis that IMDQ-PEG-CHOL is a potent adjuvant that focuses its activity to draining lymphoid tissue, combined with a promising safety profile.

**Figure 3.**
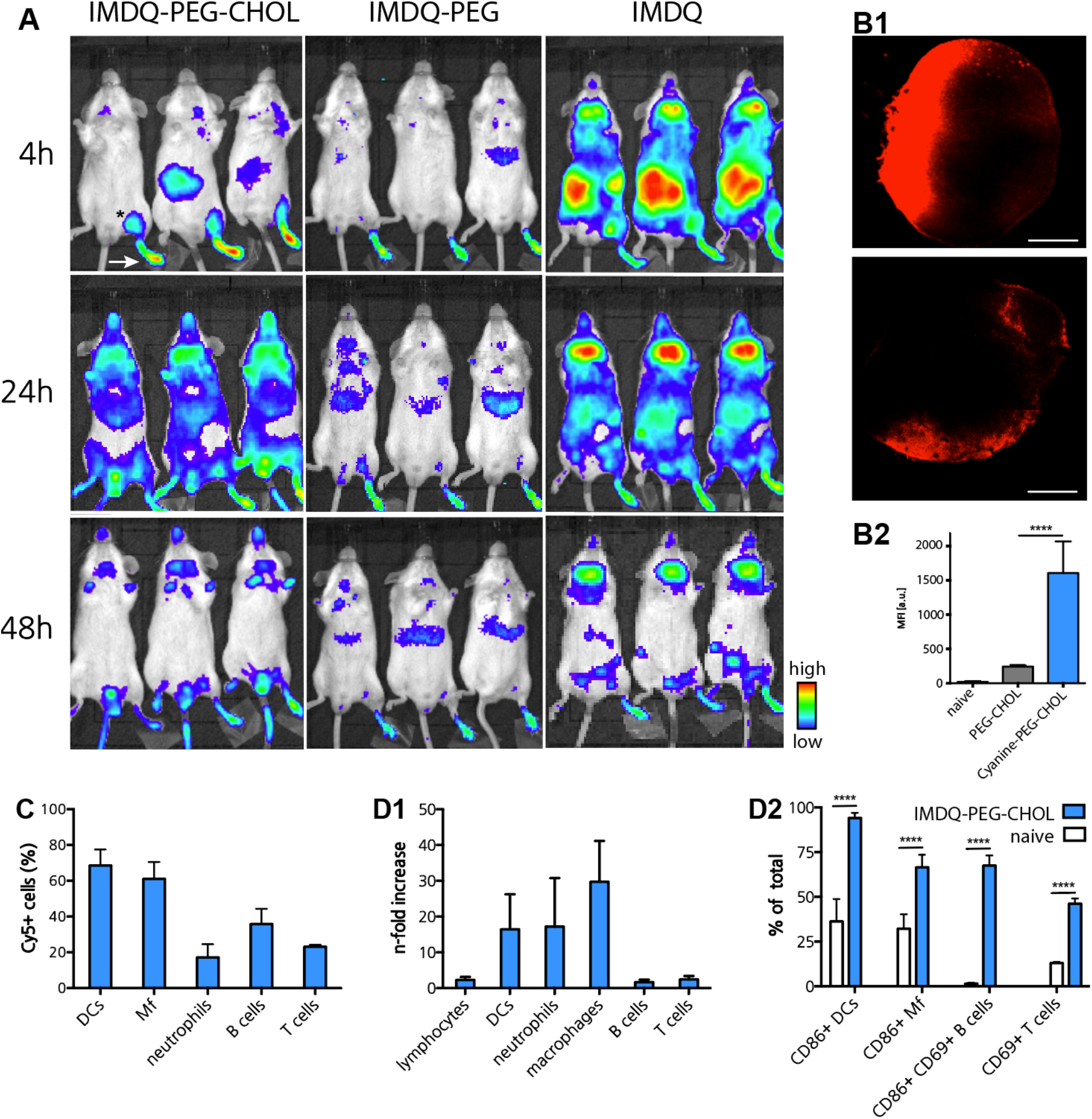
**(A)** Bioluminescence images of luciferase reporter mice (IFNβ+/Δβ-luc); images taken 4, 24 and 48 h post footpad injection of IMDQ-PEG-CHOL, IMDQ-PEG and native IMDQ. **(B1)** Confocal microscopy images of lymph node tissue sections 48 h post subcutaneous injection of Cyanine5-PEG-CHOL, respectively Cyanine5-PEG, into the footpad of mice. Scale bar represents 100 micron. **(B2)** Flow cytometry analysis of the draining popliteal lymph node 48 h post subcutaneous injection of Cyanine5-PEG-CHOL, respectively Cyanine5-PEG into the footpad of mice. (n=3, mean + SD; Student’s t-test: ****:p<0.0001) **(C)** Translocation of Cyanine5-PEG-CHOL to the draining popliteal lymph node analyzed 24 h post injection into the footpad, measured by flow cytometry. (n=6, mean + SD) **(D)** Flow cytometry analysis of the innate immune response in the draining popliteal lymph node 24 h post injection of IMDQ-PEG-CHOL into the footpad **(D1)** Relative increase in innate immune cell subsets, B and T cell numbers relative to a naïve control and **(D2)** maturation/activation of innate immune cell subsets, B and T cells (n=6, mean + SD; Student’s t-test: ****:p<0.0001).

### IMDQ-PEG-CHOL induces a balanced neutralizing antibody response to influenza virus vaccine

We evaluated the potential of IMDQ-PEG-CHOL to adjuvant a licensed vaccine, i.e. the quadrivalent influenza vaccine (QIV), in a well-established preclinical vaccination-infection model. Hereto we vaccinated BALB/c mice with each of 1.5 μg of QIV with or without 100 μg of IMDQ-PEG-CHOL or PEG-CHOL as a control. The study protocol is outlined in Figure 4A. Six mice in each group received a total of 100 μl vaccine-adjuvant mixture, intramuscularly, divided over both hind legs and the blood was collected 3 weeks post vaccination, followed by serological assays. For detection of influenza-specific antibodies induced by the QIV vaccines, we used vaccine antigen to coat enzyme-linked immunosorbent assay (ELISA) plates. The total IgG antibody titers in mice which received QIV only or QIV+ PEG-CHOL were very low as compared to the mice vaccinated with QIV+ IMDQ-PEG-CHOL (also shown as area under curve (AUC) in Supplementary Figure S4). Unadjuvanted QIV and PEG-CHOL admixed QIV resulted mainly in vaccine-specific IgG1 antibodies, whereas IMDQ-PEG-CHOL resulted in a balanced IgG1/IgG2a response as shown in Figure 4B. Interestingly, QIV + PEG-CHOL resulted in even lower antibody responses than QIV alone, an observation we confirmed in an independent vaccination experiment (data not shown). Four out of six mice that received QIV + IMDQ-PEG-CHOL could efficiently inhibit hemagglutination of chicken red blood cells (RBC) *in vitro*, by A/Singapore/gp1908/2015 IVR-180, the H1N1 virus component in QIV, with hemagglutination inhibition (HI) titers outperforming those of the other immunized groups (Figure 4C).

**Figure 4.**
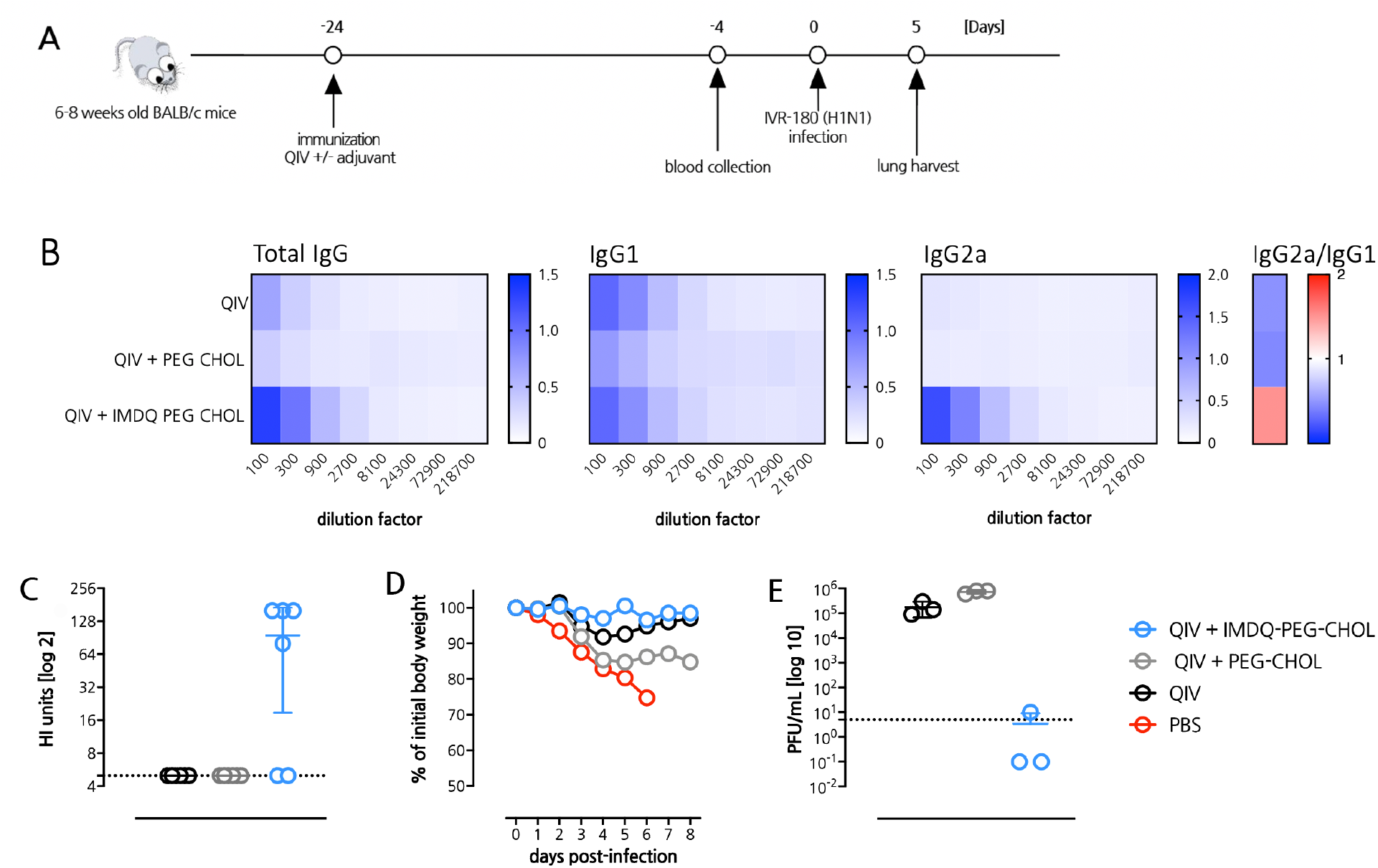
IMDQ-PEG-CHOL induces a balanced neutralizing antibody response to IVR-180 [Influenza A /Singapore /gp1908 /2015 (H1N1)] infection. **(A)** Outline of the QIV immunization and influenza virus challenge study. **(B)** Vaccine-specific ELISA titers (titers are expressed on the X axis as the reciprocal of the dilution factor) for total IgG, IgG1 and IgG2a and IgG2a/IgG1 ratio (based on the AUC (OD at 450nm) curve of the individual serum samples) in mice sera collected 3 weeks post-vaccination. **(C)** Control versus immunized sera analyzed for HI titers by hemagglutination inhibition assay, using 4 haemagglutination units of IVR-180 virus. **(D)** Body weight loss of mice reported as percentage of initial body weight after challenge with 100 LD_50_ (18000 PFU) of IVR-180 virus. **(E)** Viral lung titers after challenge with 100 LD_50_ (18000 PFU) of IVR-180 virus. Data are represented as plaque-forming-unit (PFU)/mL (geometric mean + SD). Lungs were harvested on day-5 post infection with IVR-180 virus.

Next, the immunized mice were challenged with a hundred-fold half-lethal dose (LD_50_) of IVR-180 (H1N1) virus to examine the magnitude of protection against viral infection *in vivo*. The lungs were harvested from three mice in each group 5 days post challenge to determine lung virus titers. Consistent with the ELISA and HI data, mice immunized with QIV + IMDQ-PEG-CHOL exhibited best reduction in viral lung titers as evidenced by an almost negligible number of plaques when compared to other groups (Figure 4E). This was also reflected in the optimal protection from body weight loss of QIV + IMDQ-PEG-CHOL mice after viral challenge (Figure 4D). In conclusion, our novel adjuvant IMDQ-PEG-CHOL was able to offer excellent control of viral infection and therefore, in combination with the right antigen, might also hold promise to confer protective immunity against other respiratory viruses such as SARS-CoV-2.

### IMDQ-PEG-CHOL induces a balanced neutralizing antibody response to SARS-CoV-2 immunization

We next investigated the potential of IMDQ-PEG-CHOL to adjuvant recombinant SARS-CoV-2 S protein. For this purpose, BALB/c mice were immunized intramuscularly with 6 μg of recombinant trimeric spike protein either unadjuvanted or adjuvanted with IMDQ-PEG-CHOL or with equivalent amounts of MF59-like water-in-oil vaccine AddaVax as a control established vaccine adjuvant. The recombinant vaccine consisted of the ectodomain of the SARS-CoV-2 spike protein from which the polybasic cleavage site was removed. Stabilizing prolines were added at positions 986 and 987 and trimerization was promoted by fusion to a T4 trimerization domain (see methods section for more details). The study protocol is outlined in Figure 5A. Serum was collected after 21 days post immunization and analyzed for Spike protein specific IgG titers. Whereas non-immunized mice evidently did not show any detectable Spike protein-specific titers in their sera, immunization with S protein induced Spike protein-specific titers in all groups (Figure 5B), also shown as area under the curve (AUC) titers in Supplementary Figure 5B. The total S protein-specific IgG titers were found to be the highest in the IMDQ-PEG-CHOL adjuvanted group. Additionally, Spike protein + IMDQ-PEG-CHOL immunization resulted in a higher IgG2a/IgG1 ratio, suggesting a more potent Th1 immune response and more efficient class switching towards IgG2a as compared to spike protein only or spike protein + AddaVax immunization.

**Figure 5.**
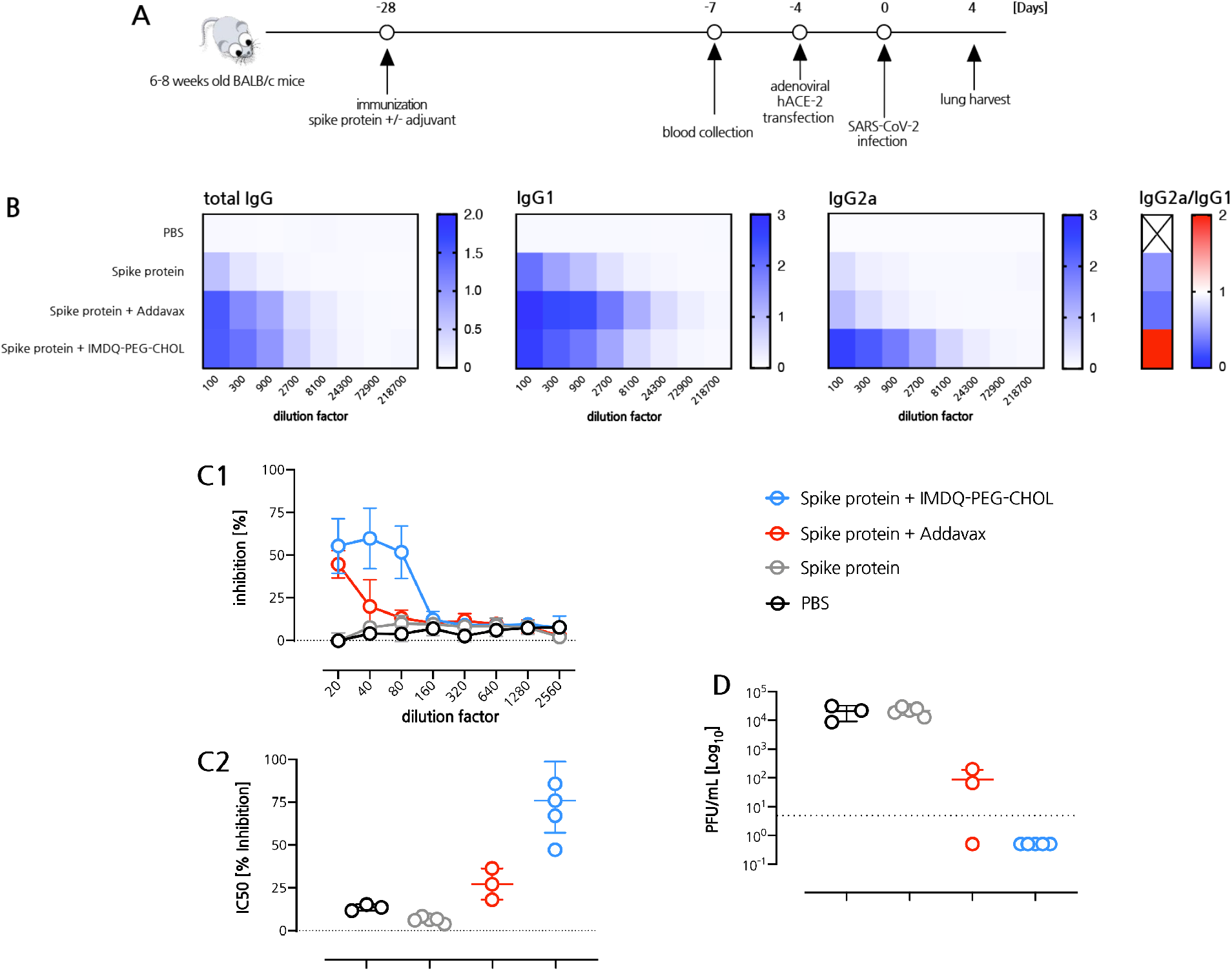
IMDQ-PEG-CHOL induces a balanced neutralizing antibody response to SARS-CoV-2 infection. **(A)** Outline of the Spike protein vaccination and SARS-CoV-2 challenge. **(B)** ELISA titers (titers are expressed on the X axis as the reciprocal of the dilution factor) for total IgG, IgG1 and IgG2a and IgG2a/IgG1 ratio (based on the AUC (OD at 450nm) curve of the individual serum samples) in mice sera collected 3 weeks post-vaccination. **(C)** Control versus vaccinated sera examined for presence of virus-neutralizing antibodies by microneutralization assay, using 100 tissue culture infectious dose 50 (TCID50) of SARS-CoV-2 virus. The outcome is represented as a percentage inhibition of viral growth in (C1) and as the half maximal inhibitory concentration IC_50_ calculated by a non-linear regression analysis of percentage inhibition curve in (C2). **(D)** Viral lung titers represented as Plaque-forming-unit (PFU)/mL (geometric mean with geometric SD). The Ad5-hACE2 transduced mice were challenged with 5×10^4^ PFU of SARS-CoV-2 and the lungs were harvested on day-4 post infection.

Next, the sera from vaccinated mice were used to test the ability to inhibit SARS-CoV-2 infection *in vitro*. Although immunization with non-adjuvanted S protein was able to induce some IgG titers, it was found ineffective in neutralizing viral infection of Vero E6 cells *in vitro* in a microneutralization assay (Figure 5C1-2). By contrast, serum of mice immunized with spike protein + IMDQ-PEG-CHOL was able to neutralize >50% of SARS-CoV-2 virus infection in this assay, which was also significantly higher than the serum of mice immunized with spike protein + AddaVax.

### IMDQ-PEG-CHOL induces protective immunity against SARS-CoV-2 in a mouse model

Finally, we investigated to what extent S protein + IMDQ-PEG-CHOL immunization is able to confer protection against a SARS-CoV-2 viral challenge. As mice do not express the hACE-2 receptor that is needed for the virus to infect the host, we first transduced immunized mice with an adenoviral vector encoding for hACE-2 by intranasal installation. Four days later, mice we challenged with SARS-CoV-2 virus and again 4 days later, lungs were harvested, and the residual viral infection was quantified by a plaque assay. Interestingly, whereas non-adjuvanted Spike protein could not confer any protection, relative to the non-immunized group, Spike protein + IMDQ-PEG-CHOL vaccination conferred sterilizing immunity against the SARS-CoV-2 infection with plaque numbers below the detection limit (Figure 5D), and performed significantly better than spike protein + AddaVax immunization, which correlates with the higher microneutralization titers observed in the Spike protein + IMDQ-PEG-CHOL group (Figure 5C1-2).

## Conclusions

In summary, we have shown in this work that IMDQ-PEG-CHOL is a potent adjuvant with enhanced safety profile that induced innate immune activation in lymphoid tissue upon local administration. Whereas IMDQ in soluble, unformulated form, rapidly enters systemic circulation, conjugation to a lipid-polymer amphiphile prevents the latter while promoting translocation to the draining lymph node, likely through binding to albumin in the interstitial flow. In mouse vaccination/challenge models for influenza and SARS-CoV-2, we have demonstrated that a single immunization with QIV or S protein adjuvanted with IMDQ-PEG-CHOL induced robust Th1 skewed antibody responses, as evidenced by higher IgG2a/IgG1 ratios. Importantly, vaccination with QIV and S protein adjuvanted with IMDQ-PEG-CHOL resulted in virus-specific neutralizing antibodies and control of viral infection after challenge with influenza and SARS-CoV-2 viruses, respectively. Since IgG2a subclass is known to engage Fcγ receptors that are involved in antiviral protection mechanisms like antibody-mediated cellular cytotoxicity and phagocytosis, IMDQ-PEG-CHOL might be beneficial to promote such responses. Whereas we are aware of the limitations of the present studies, we do believe that the overarching message that single vaccination with a properly adjuvanted recombinant S protein based vaccine is able to induce protective immunity in a mouse model, is of great relevance with regard to broadening the arsenal of emerging COVID-19 vaccines. Safe adjuvants like IMDQ-PEG-CHOL described in our study may help enhance vaccine efficiency if deemed necessary from ongoing clinical trials, similar to the recent development of MF59-adjuvanted influenza virus vaccines for the elderly. Use of efficient adjuvants may also reduce the amount of vaccine needed to induce a protective immune response, which is important in case there is vaccine shortage, as is the case during this COVID19 pandemic.

## Materials and Methods

### Materials

All chemicals for synthesis were purchased from Sigma-Aldrich or TCI, unless noted otherwise.

### Mice, Cell lines and reagents

6-8 weeks old female BALB/c mice were obtained from Charles River Laboratories, MA and were housed in a specified pathogen-free facility at Icahn school of medicine at Mount Sinai, with food and water ad libitum, adhering to the guidelines from Institutional Animal Care and Use Committee.

Madin-Darby Canine Kidney Cells (MDCK, ATCC-CCL 34) and Vero-E6 (ATTC-CRL 1586, clone E6) cells are routinely cultured in the laboratory. Both cells were maintained in Dulbecco’s Modified Eagle’s Medium supplemented with 10% Fetal bovine serum (FBS) and additionally with 1% non-essential amino acids for Vero-E6 cells. DC2.4 mouse dendritic cells were cultured in RPMI-glutamax supplemented with 10 % fetal bovine serum (FBS), antibiotics (50 units/mL penicillin and 50 *μ*g/mL streptomycin) and 1 mM sodium pyruvate. Murine RAW blue 264.7 macrophages were cultured in DMEM medium supplemented with 10 % heat-inactivated FBS, antibiotics (50 units/mL penicillin and 50 *μ*g/mL streptomycin), 2 mM L-glutamine and 0.01 % Zeocin. Cells were incubated at 37 °C in a controlled and sterile environment of 95 % relative humidity and 5 % CO_2_.

### Vaccines, Blood Collection and Serology

#### Quadrivalent inactivated influenza virus vaccine (QIV)

was the human Seqirus vaccine (2018-2019 formula) containing the antigens of the following influenza virus strains: A/Singapore/GP1908/2015 IVR-180 (H1N1), A/North Carolina/04/2016 (H3N2), B/Iowa/06/2017 and B/Singapore/INFTT-16-0610/2016. Vaccine was obtained from BEI resources and mixed with adjuvant as described below. Vaccine was injected once via the intramuscular route with a BD 300 μL insulin syringe in the hamstring muscles of the both hind legs (50 μL/leg). The administered vaccine dose corresponds to 1.5 μg of each hemagglutinin type in the vaccine per mouse.

Blood was collected twenty days post vaccination via submandibular bleeding and serum was prepared by allowing the blood to clot at room temperature. Anti-HA antibody responses were measured by enzyme linked immunosorbent assay (ELISA) and hemagglutination inhibition (HI) assay. For quantification of HA-specific total IgG levels by ELISA, 96 well NUNC Maxisorp plates were coated with QIV (2 μg/ml HA equivalent for each HA) in bicarbonate buffer at 4 °C overnight. After washing and blocking with 4% milk for 1 h at room temperature, serum samples 3-fold diluted starting at 1/100 in PBS with 0.05% Tween20 were allowed to bind ELISA antigen for 1.5 h at room temperature. Plates were washed three times with PBS (0.05% Tween20) and incubated with sheep-derived anti-mouse total IgG (GE Healthcare, Amersham, UK), IgG1 or IgG2a (Invitrogen) serum conjugated to horse-radish peroxidase. After a final washing step, tetramethylbenzidine (TMB) substrate (Sigma-Aldrich, San Diego, CA, USA) was used to estimate levels of HA-specific mouse IgG by measuring the OD_450_ with the OD_650_ as a reference after stopping the colorimetric reaction with 1M H_2_SO_4_.

Hemagglutination inhibition was performed as previously described (23). Briefly, four volumes of receptor destroying enzyme (RDE, Vibrio cholera filtrate, Sigma Aldrich, San Diego, CA, USA) were added to each volume of mouse serum. After overnight incubation at 37 °C, sera were heat-inactivated at 56 °C for 30 min in sodium citrate buffer. Four hemagglutination units of IVR-180 H1N1 virus were mixed with twofold dilutions of treated sera in a final volume of 50 μL. Mixtures of virus and diluted serum were allowed to bind for 1h at room temperature before 50 μL of 0.5% chicken red blood cell suspension was added. HI titers were read after 1h incubation on ice.

#### Trimeric recombinant SARS-CoV-2 spike protein

was produced as previously described: only the ectodomain of the spike protein (GenBank: MN908947.3) was cloned into a mammalian expression plasmid and the cleavage site was removed and stabilizing prolines were added at position 986 and 987 (24–26). A hexa-histidine tag as well as a T4 foldon trimerization domain was present in the plasmid for ease of purification. The spike protein was expressed in 293F cells, using the ExpiFectamine 293 Transfection Kit (Thermo Fisher). Supernatant was collected on day 3 post transfection and Ni-NTA agarose (Qiagen) was used to purify the protein. This protocol has been described in much greater detail earlier (26). Vaccine (6 μg/mouse) was mixed with adjuvant as described below and injected once via the intramuscular route with a BD 300 μL insulin syringe in the hamstring muscles of both hind legs (50 μL/leg).

Anti-SARS-CoV-2 spike protein ELISA was performed to estimate spike-specific antibody responses upon vaccination. Briefly, maxisorp Nunc 96-well microtiter plates were coated with 50 μl per well of recombinant spike protein, diluted to a concentration of 2 μg/ml in carbonate/bicarbonate buffer and incubated overnight at 4°C. Three-fold serially diluted serum samples, starting from 1:100, were added to the antigen-coated plates followed by overnight incubation at 4°C. The plates were then washed in 1X PBS + 0.01% Tween20 and again incubated with appropriate horse-radish peroxidase (HRP)-conjugated secondary antibodies targeting total IgG, IgG1 or IgG2a antibodies (GE Healthcare). The plates were washed and developed with 50 μl of TMB substrate per well until blue color appeared. The reaction was terminated with 50 μl 1M H_2_SO_4_ and the absorbance was measured at 450nm with 650 nm as a reference.

#### In vitro microneutralization assay

To measure the neutralizing potential of SARS-CoV-2 vaccine-induced sera, an *in vitro* microneutralization assay was performed similar to the protocol described in (27). Briefly, the Spike ± adjuvant-vaccinated mice sera were inactivated at 56°C for 30 min. Serum samples were serially diluted 2-fold starting from 1:10 dilution in infection medium (DMEM+ 2% FBS+ 1X non-essential amino acids). The samples were incubated with 100 tissue culture infective dose 50 (TCID50) which equals 40 plaque forming units (PFU) of SARS-CoV2 virus for 1 hour in an incubator at 37°C, 5% CO_2_ and then transfer on pre-seeded Vero-E6 cells in 96-well cell-culture plates. The plates were incubated at 37°C for 48 hours and fixed in 4% formaldehyde. The cells were washed with 1XPBS and blocked in 5% milk in 1XPBS+ 0.1% Tween20 for 1 hour at room temperature. After blocking, the cells were permeabilized with 0.1% TritonX100, washed and incubated with anti-SARS-CoV-2-nucleoprotein and anti-SARS-CoV-2-Spike monoclonal antibodies, mixed in 1:1 ratio, for 1.5 hours at room temperature. The cells were washed again and incubated with HRP-conjugated anti-mouse IgG secondary antibody for 1 hour at room temperature followed by a brief PBS wash. Finally, 50 μl tetramethyl benzidine (TMB) substrate was added and incubated until blue color appeared and the reaction was terminated with 50 μl 1M H_2_SO_4_. Absorbance at 450nm was recorded and percentage inhibition calculated. Anti-mouse SARS-CoV-2-nucleoprotein and anti-mouse SARS-CoV-2-Spike antibodies were obtained from the Center for Therapeutic Antibody Development at the Icahn School of Medicine at Mount Sinai, New Y ork.

#### Adjuvants

AddaVax was purchased from Invivogen and mixed at a 3:1 ratio vaccine:AddaVax per the manufacturer’s recommendation. IMDQ adjuvants were mixed with vaccine at an equivalent of 10 μg core IMDQ (100 μg of IMDQ-PEG-CHOL, see below) per mouse.

### Viruses and Infection

QIV vaccinated mice were infected 24 days post vaccination with 100 lethal dose 50 (18,000 PFU) of egg-grown influenza IVR-180 H1N1 virus, a vaccine strain that contains the surface antigens of influenza A/Singapore/gp1908/2015 (H1N1) virus. Morbidity and mortality were monitored for eight days. A group of age-matched naïve animals was added to the experiment to confirm the dose of virus was lethal to unvaccinated animals.

In order to make SARS-CoV-2 Spike-vaccinated BALB/c mice susceptible to challenge with wild type SARS-CoV-2 virus, airway expression of human ACE-2, the receptor for SARS-CoV-2, was obtained by intranasal transduction of mice with 2.5×10^8^ PFU of adenovirus expressing h-ACE-2 (Ad5-hACE2), 4.5-weeks post-vaccination as described in (28). Five days after transduction with Ad5-hACE2, mice were challenged with 5×10^4^ PFU of SARS-CoV2 isolate USA-WA1/2020 (BEI resources; NR-52281) per mice. Body weights were recorded to assess the morbidity during the days post challenge.

### Lung Virus Titration

Plaque assays were performed to quantify and compare the lung viral titers in vaccinated versus unvaccinated mice. As described previously (23), whole lungs were harvested from the mice and homogenized in 1 ml 1XPBS. After brief centrifugation, the tissue debris was discarded and the supernatant was 10-fold serially diluted starting from 1:10 dilution. For IVR-180, MDCK cells were incubated with the lung homogenate dilutions for 1 hour at 37°C, 5% CO_2_ and then overlaid with a mixture of 2% oxoid agar and 2X minimal essential medium (MEM) supplemented with 1% diethyl-aminoethyl (DEAE)-dextran and 1 μg/ml tosylamide-2-phenylethyl chloro-methyl ketone (TPCK)-treated trypsin. After 48 hours of incubation at 37°C, 5% CO_2_, the plates were fixed in 4% formaldehyde and immune-stained with IVR-180-post-challenge polyclonal serum. Similarly, For SARS-CoV-2, pre-seeded Vero-E6 cells were incubated with diluted lung homogenates for 1 hour at room temperature and then overlayed with a 1ml mixture of 2% oxoid agar and 2X MEM supplemented with 2% FBS. After 72 hours of incubation at 37°C, 5% CO_2_, the plates were fixed in 4% formaldehyde, followed by immune-staining of infected cells with anti-mouse SARS-CoV-2 nucleoprotein and anti-mouse SARS-CoV-2 spike monoclonal antibodies. After incubation in primary antibodies, HRP-conjugated anti-mouse secondary antibody was added for 1 hour. Finally, the plaques were developed with TrueBlue substrate (KPL-Seracare). The final viral titers were calculated in terms of plaque forming units (PFU)/ml.

### IMDQ-PEG-CHOL synthesis

IMDQ was synthesized according to literature (18).

#### Synthesis of Cholesterylamine

First, the alcoholic hydroxyl group of cholesterol was transformed to an azide through a Mitsunobu reaction in the presence of diphenylphosphoryl azide (DPPA). Cholesterol (2.0 g, 5.17 mmol) was dissolved in a round bottom flask equipped with a stirring bar containing anhydrous THF (20 mL). Triphenylphosphine (PPh_3_, 1.63 g, 6.21 mmol) and diisopropyl azodicarboxylate (DIAD, 1.22 mL, 6.21 mmol) was added to the round bottom flask. Upon addition of DIAD, the reaction mixture developed a yellow colour. After 10 minutes, DPPA (1.34 mL, 6.21 mmol) was added and the mixture stirred overnight at room temperature under inert atmosphere. The reaction mixture was reduced under vacuum and further purified by column chromatography (cyclohexane), to yield a purified withe powder (yield = 60 %). The resulting product cholesteryl-N_3_ was analysed by ^1^H-NMR and ATR-IR.

Next, a Staudinger reduction was executed to reduce the azide group to a primary amine function with PPh_3_. The obtained cholesteryl-N_3_ (400 mg, 0.97 mmol) was transferred into a round bottom flask containing a stirring bar and dissolved in anhydrous THF (2.0 mL) under inert atmosphere. A solution of PPh_3_ (2.55 g, 9.72 mmol) in dry THF (5.0 mL) was added. After 30 min, 2 mL water was added and the reaction mixture stirred overnight at room temperature equipped with a balloon to trap the released nitrogen gas. The reaction mixture was diluted extensively with toluene before being reduced under vacuum by 50 °C. The crude product was purified by column chromatography using a gradient (from 95:5 DCM:MeOH to 95:5 DCM:MeOH + 1% ammonium hydroxide), yielded a white powder which was characterized by ^1^H-NMR, ATR-IR and MS(yield = 98 %). ESI-MS (Figure SX): m/z [M+H]^+^ = 386.37 (theoretical); found = 386.363

#### Synthesis of PEG-CHOL

A round bottom flask containing a stirring bar was loaded with Boc-NH-PEG-COOH (300 mg, 0.098 mmol) and dissolved in anhydrous *N,N*-dimethylformamide (DMF, 3 mL) under inert atmosphere. 1 - [Bis(dimethylamino)methylene]-1H-1,2,3-triazolo[4,5-b]pyridinium 3-oxid hexafluorophosphate (HATU) (40.84 mg, 0.107 mmol) was added to the stirring solution followed by 20.4 *μ*L of triethylamine (TEA, 0.146 mmol). After 5 min, a solution of cholesterylamine (56.4 mg, 0.146 mmol) in dry chloroform (1.5 mL) was added and stirred at room temperature for 2 h, yielding a slightly yellow reaction mixture. The reaction mixture was co-evaporated with a large excess of toluene to remove DMF. Subsequently, the crude product was purified by threefold precipitation in ice cold diethyl ether followed by column chromatography (90:10 DCM: MeOH) (yield = 86 %). The white PEG-CHOL-NH Boc powder was analyzed by ^1^H-NMR, DMAc-SEC and MALDI-ToF.

Subsequently the Boc-group was removed by dissolving Boc-NH-PEG-CHOL (200 mg, 0.058 mmol) in 2 mL of dichloromethane (DCM) in a round bottom flask equipped with a stirring bar. An equal amount of trifluoroacetic acid (TFA, 2 mL) was added and the solution was stirred for 2 h opened to ambient air and temperature. Prior to concentration under vacuum, a large excess of toluene was added to the reaction mixture. Finally, the product was transferred to dialysis membranes and dialyzed against 0.1 % v/v ammonium hydroxide solution in demineralized water for multiple days and one day against demineralized water. After lyophilization, the white fluffy powder NH_2_-PEG-CHOL was characterized by ^1^ H-NMR and MALDI-ToF.

#### Synthesis of **linker 1**

In a round bottom flask equipped with a stirring bar, p-nitrobenzyl chloroformate (2.02 g, 10 mmol) was dissolved in anhydrous DCM and cooled on ice. A mixture of diethylene glycol (424.5 mg, 4 mmol) and TEA (1.67 mL, 12 mmol) in 10 mL anhydrous DCM were added dropwise and stirred for an additional 30 minutes on ice. After 2 h on room temperature, the reaction mixture was concentrated under vacuum, dissolved in EtOAc and filtered. After evaporation of the solvent under reduced pressure, the linker was analyzed by ^1^H-NMR and MS.

#### Synthesis of IMDQ-PEG-CHOL

First, **linker 1** (65.3 mg, 0.15 mmol) was dissolved in 2.5 mL anhydrous DCM under inert atmosphere. After addition of 2 equivalent of dry TEA (8.35 *μ*L, 0.06 mmol), NH_2_-PEG-CHOL (100 mg in 2 mL anhydrous DCM, 0.03 mmol) was added dropwise under stirring and a distinct yellow color appeared indicating the release of p-nitrophenol. After overnight reaction, purification was performed by double precipitation into a mixture of ice-cold hexane:acetone (80:20). The resulting **intermediate 1** was dried under vacuum and analyzed by NMR and SEC with DMAc as mobile phase.

In the second step, **intermediate 1** (50 mg, 0.014 mmol) and IMDQ (9.5 mg, 0.22 mmol) were weighed into a round bottom flask with stirring bar. The compounds were dissolved in 2.3 mL anhydrous 1,4-dioxane and anhydrous TEA (9.76 *μ*L, 0.07 mmol) was added to solution under vigorous stirring. After 3 h at room temperature, a few drops of dry methanol were added and the reaction was stirred overnight. Next, the solution was transferred into a dialysis membrane (1 kDa) and dialyzed for multiple days against demineralized water. After freeze drying, the with fluffy*IMDQ-PEG-CHOL* powder was characterized by ^1^H-NMR, HPLC and MALDI-ToF. HPLC conditions to verify absence of freely soluble IMDQ: LiChroCart® C18 column 250-4, mobile phase H_2_O/ ACN 65:35 with 0.1% TFA, flow rate at 1 mL/min and detection at 250nm. Note that IMDQ-PEG was synthesized in similar fashion, but omitting the conjugation of cholesterylamine.

#### Synthesis of Cyanine5-PEG-CHOL

NH_2_-PEG-CHOL (25 mg, 7.49 *μ*mol) was weighted into a Schlenk tube equipped with a magnetic stirring bar and dissolved in 2.5 mL dry DMSO under inert atmosphere. Then, 0.21 mL of cyanine5 N-hydroxysuccinimide ester (stock solution of 25 mg/mL in anhydrous DMSO, 7.86 μmol) and anhydrous TEA (5.2 μL,37.42 μmol) were added to the Schlenk tube and further stirred overnight at room temperature. After dialyzed for three days against demineralized water, Cyanine5-PEG-CHOL was isolated as a fluffy blueish powder after lyophilization.

### Biolayer interferometry analysis

Bovine serum albumin (BSA) was biotinylated by reacting it with 5:1 excess of biotin-NHS followed by dialysis and lyophilization. Hydrated streptavidin sensors were dipped in PBS to record a baseline for 60 seconds, followed by dipping into biotinylated BSA (12.5 nM, 66.5 kDa) in PBS for 300 s, and dipping for 30 s in PBS for washing. Next, a second baseline was recorded by dipping in fresh PBS for 120 s. Association of IMDQ-PEG-CHOL was measured by dipping into a solution of IMDQ-PEG-CHOL in PBS for 600 s. Note that the experiment was ran in parallel for different concentrations of IMDQ-PEG-CHOL. Dissociation of IMDQ-PEG-CHOL was recorded by dipping in PBs for 600 s. The experiment was performed in a black flat bottom 96 well plate set at 30 °C by 1000 rpm, using an Octet RED96 model (Pall Fortébio). Data processing was done by the FortéBio software package.

### Cell cytotoxicity (MTT) assay

DC 2.4 cells were plated seeded in 96-well plates at a density of 8 000 cells per well in 200 μL culture medium. 50 μL of sample (dilution series in PBS, ranging from 10^−4^ mg/mL to 0.5 mg/mL), PBS (negative control, 100 % viability) and DMSO (positive control, 0 % viability) were added to the wells. After 72 h, the medium was aspirated and cells were washed with 200 μL PBS followed by addition of 100 μL of diluted MTT stock solution. After 1h, the solution was removed and the formed formazan crystals were dissolved in 50 μL DMSO. Quantification was done by measuring the absorbance at 590 nm using a microplate reader. Note, 3-(4,5-dimethylthiazol-2-yl)-2,5-diphenyltetrazolium bromide (MTT, 50 mg) was dissolved in 10 mL sterile PBS, filtrated (membrane 0.22 μm) and 1/5 diluted in culture medium prior to use in this assay.

### Confocal microscopy

DC 2.4 cells were seeded in Willco-Dish glass bottom at a concentration of 10 000 cells in 180 μL culture medium and allowed to adhere overnight. Cells were pulsed overnight with 20 μL of a 1 mg/mL Cyanine5-PEG-CHOL or Cynanine5-PEG solution in PBS. Next, the culture medium was aspirated and cells were fixated with 4% paraformaldehyde (PFA) for 30 min followed by washing with PBS and confocal imaging using a Leica DMI6000B microscope (63x 1.40 NA objective) coupled to an AndorDSD2 confocal scanner and a Zyla5.5 CMOS camera. Image processing was done using the ImageJ software package.

### Flow cytometry analysis of in vitro IMDQ-PEG-CHOL association

DC 2.4 cells were seeded out in 24-well plate at a concentration of 200 000 cells per well in 450 μL of culture medium. Cells were pulsed with samples and incubated overnight at 37 °C. Afterwards, the supernatants were removed, cells were washed with PBS and detached with cell dissociation buffer (0.5 mL, 15 min, 37 °C). The content of the wells was transferred to an Eppendorf and centrifuged (5 min, 300 G, 4 °C). After aspiration of the supernatant, the cell pellets were resuspended in PBS and analyzed using a BD Accuri Flow Cytometer. Data were processed using the FlowJo software package.

### RAW blue innate immune activation assay

RAW blue 264.7 macrophages were seeded in flat-bottom 96 well plate at a density of 70.000 cells per well, suspended in 180 μL culture medium and pulsed with 20 μL of sample for 24h at 37 °C at different concentrations of IMDQ-PEG-CHOL, IMDQ-PEG, IMDQ and PBS. Subsequently, 50 μL of supernatant was transfer to a new flat-bottom 96 well plate followed by addition of 150 μL of QUANTI-Blue™ reagent solution, prepared according to the manufacturer’s instruction (Invivogen). After 30 minutes at 37 °C, the SEAP levels were determined by UV-Vis spectrophotometry at 620 nm using a microplate reader. Note, the colorimetric quantification of the samples was obtained relative to the negative control and each concentration was performed in fivefold.

### Analysis of *in vivo* lymphatic drainage

20 μL (1 mg/mL in PBS) of Cyanine5-PEG-CHOL or Cyanine5-PEG were injected into the footpad of female C57BL/6 WT mice. Two mice were used per group and injected in both footpads. At the designated time point, mice where sacrificed and popliteal lymph nodes where isolated for flow cytometry and confocal imaging. A single cell suspension was prepared from the dissected popliteal lymph nodes for analysis by flow cytometry. Isolated lymph nodes were collected in ice cold PBS, dissociated through 70 μm cell strainers, washed with PBS and stained with a fixable dead/live-staining. 123count ebeads were added to determine cellularity prior to analysis by a BD FACS Quanto flow cytometer. Data were processed using the FlowJo software package.

For confocal imaging popliteal lymph nodes where frozen in OCT cryomedium (Sakura, 4583). frozen sections (8-μm) were cut by cryostat. These sections where fixed for 4 min in PFA 2% (v/v), and washed with PBS. Images were acquired on a Zeiss LSM710 confocal microscope equipped with 488-nm, 561-nm and 633-nm lasers and with a tunable two-photon laser. Confocal imaging was done using a Leica DMI6000B microscope (10x 0.70 NA objective) coupled to an AndorDSD2 confocal scanner and a Zyla5.5 CMOS camera. Image processing was done using the ImageJ software package.

### *In vivo* immune activation imaging

Luciferase reporter mice (IFNβ+/Δβ-luc) with a Balb/c background, aged 7-9 weeks, were housed in individual ventilated cages and given ad libitum access to food and water. 20 mL of IMDQ-PEG-CHOL, IMDQ-PEG or IMDQwere injected subcutaneously in the footpad (n=3) at an equivalent IMDQ dose of 2 mg. For in vivo imaging at the given time points, mice were injected subcutaneously with 200 μL D-luciferin and in vivo luminescence imaging was recorded 12 min later using the IVIS Lumina II imaging system. Local (DLN and DLN + foot pad) luminescence and full-body luminescence were quantified using the Living Image 4.4 software.

### Analysis of *in vivo* lymphocyte targeting and activation

20 μL (containing an equivalent IMDQ dose of 2mg) of IMDQ-PEG-CHOL or Cyanine5-PEG-CHOL was injected into the footpad of female C57BL/6 WT mice. At the designated time point, mice where sacrificed and popliteal lymph nodes where isolated. A single cell suspension was prepared from the dissected popliteal lymph nodes for analysis by flow cytometery. Isolated lymph nodes were collected in ice cold PBS, smashed through 70 μm cell strainers, washed with PBS and stained for 30min at 4°C with following primary labeled antibodies: CD3, CD20, CD11c, MHCII, CD86, CD40. Live dead ratio’s where determined by staining with fixable dead/live-staining and 123count ebeads were added to determine cellularity prior to acquiring them on 123count ebeads were added to determine cellularity prior to Analysis by a BD FACS Quanto flow cytometer. Data were processed using the FlowJo software package.

## Acknowledgements

This work was partly funded by CRIP (Center for Research on Influenza Pathogenesis), a NIAID funded Center of Excellence for Influenza Reserch and Surveillance (CEIRS, contract #HHSN272201400008C), by SEM-CIVIC, a NIAID funded Collaborative Influenza Vaccine Innovation Center (contract #75N93019C00051), by a supplement to NIAID contract 75N93019C00045, by NIAID grants U01AI124297 and P01AI097092, by FASTGRANT 2176, and by the generous support of the JPB Foundation, the Open Philanthropy Project (research grant 2020-215611 (5384)) and anonymous donors to AG-S. This work was supported in part by NIAID R21AI157606 (L.C). Work in the Krammer laboratory was supported by the NIAID Centers of Excellence for Influenza Research and Surveillance (CEIRS) contract HHSN272201400008C (FK, for reagent generation), Collaborative Influenza Vaccine Innovation Centers (CIVIC) contract 75N93019C00051 (FK, for reagent generation), and the generous support of the JPB foundation, the Open Philanthropy Project (#2020-215611) and other philanthropic donations. B.G.D.G. acknowledges funding from the European Research Council (ERC) under the European Union’s Horizon 2020 research and innovation program (grant N 817938). We also thank Randy Albrecht and Carles Martinez for support with the BSL3 facility and procedures at the ISMMS and Richard Cadagan for excellent technical assistance.

## Supplementary figures

**Supplementary Fig. 1:**
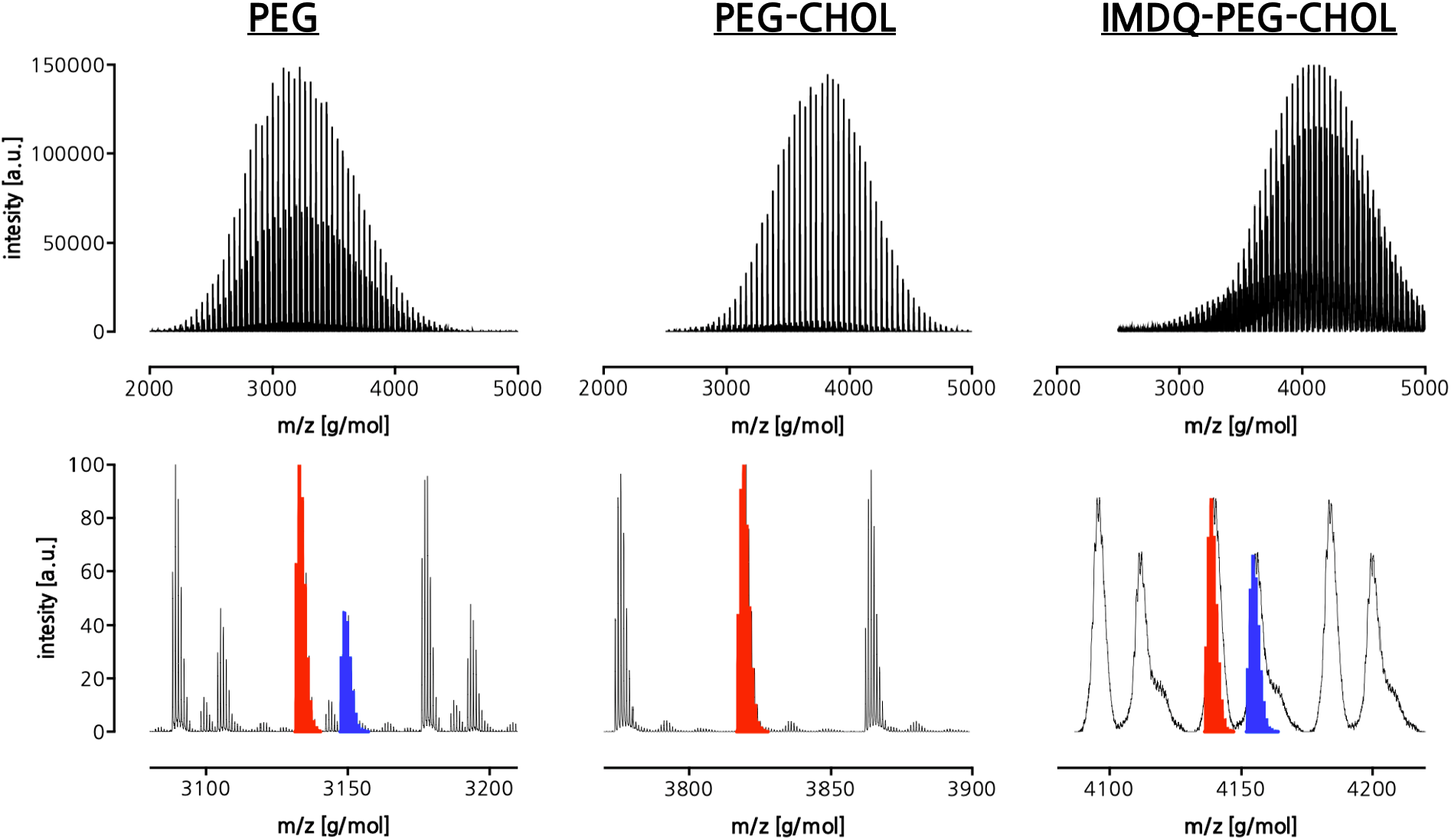
MALDI-ToF analysis of PEG, PEG-Chol and IMDQ-PEG-CHOL. The bottom row depicts a zoom highlighting the simulated sodium adduct in red and the potassium adduct in blue.

**Supplementary Fig. 2:**
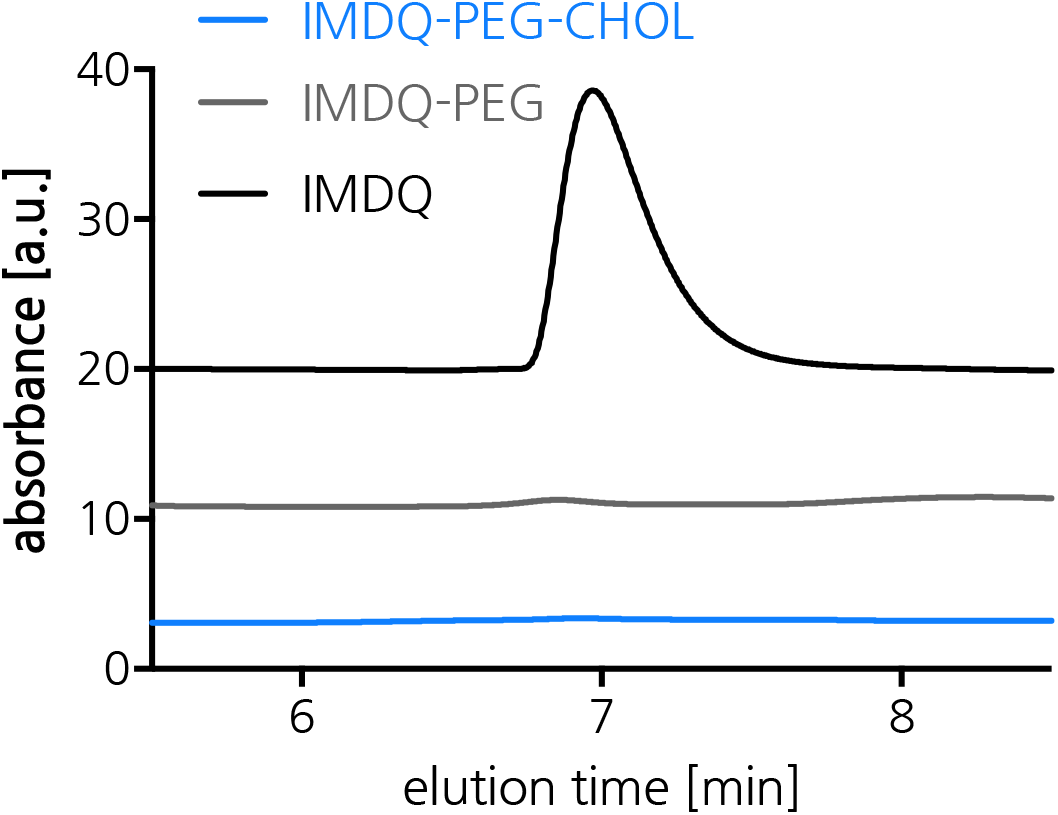
HPLC elugrams (eluens 50:50acetonitrile/water with 0.1 vol%TFA) showing absence of unmodified IMDQ in IMDQ-PEG and IMDQ-PEG-CHOL as no IMDQ peak (emerging at 7 min in the IMDQ elugram (black curve)) is observed in the IMDQ-PEG and IMDQ-PEG-CHOL elugrams (blue and grey curves).

**Supplementary Fig. 3:**
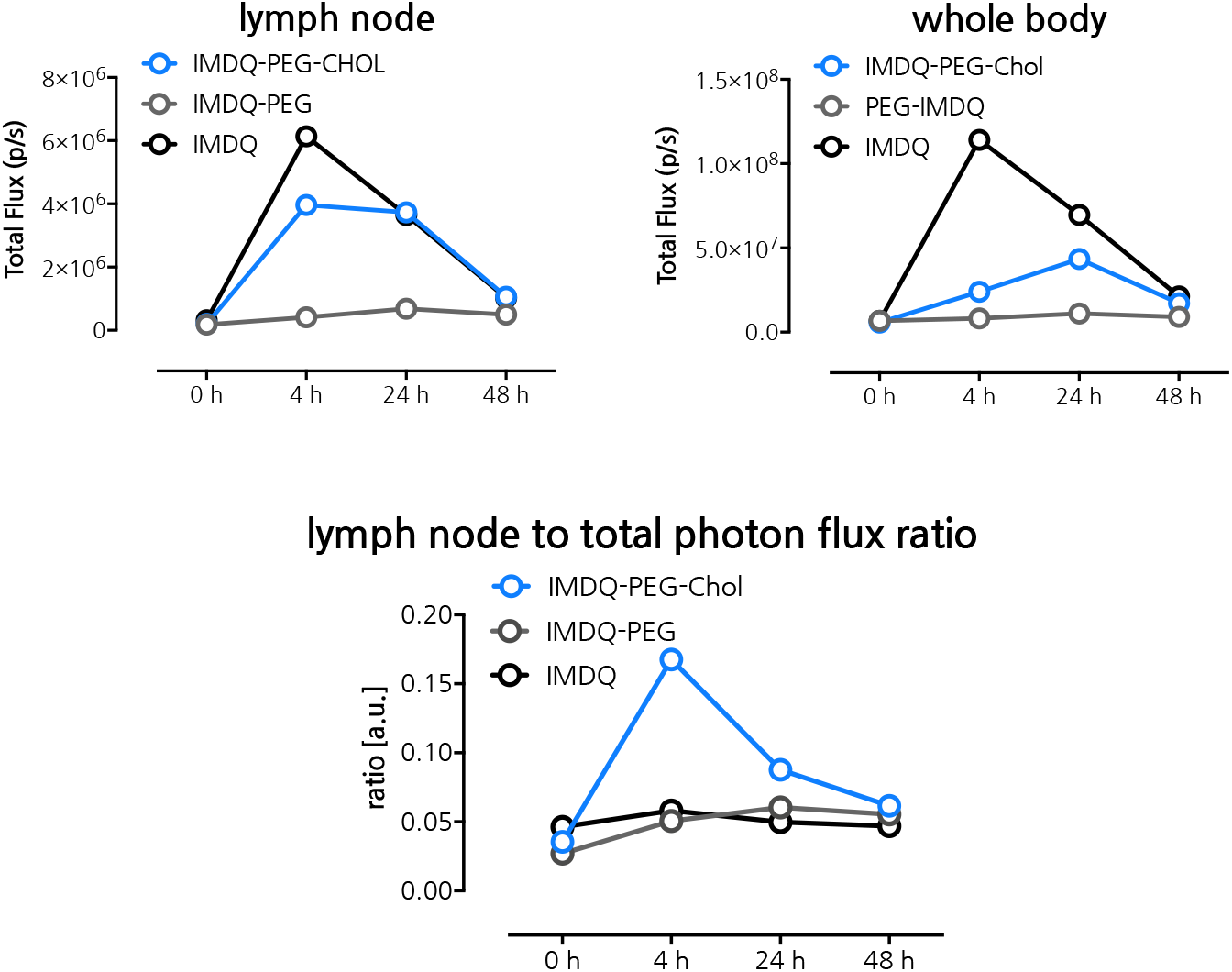
Quantification of the total photon flux from the draining popliteal lymph node and the whole body, and the ratio of both. Quantification was done based on the bioluminescence images (Figure 3A) of luciferase reporter mice (IFNβ+/Δβ-luc) recorded 4, 24 and 48 h post footpad injection of IMDQ-PEG-CHOL, IMDQ-PEG and native IMDQ

**Supplementary Fig. 4:**
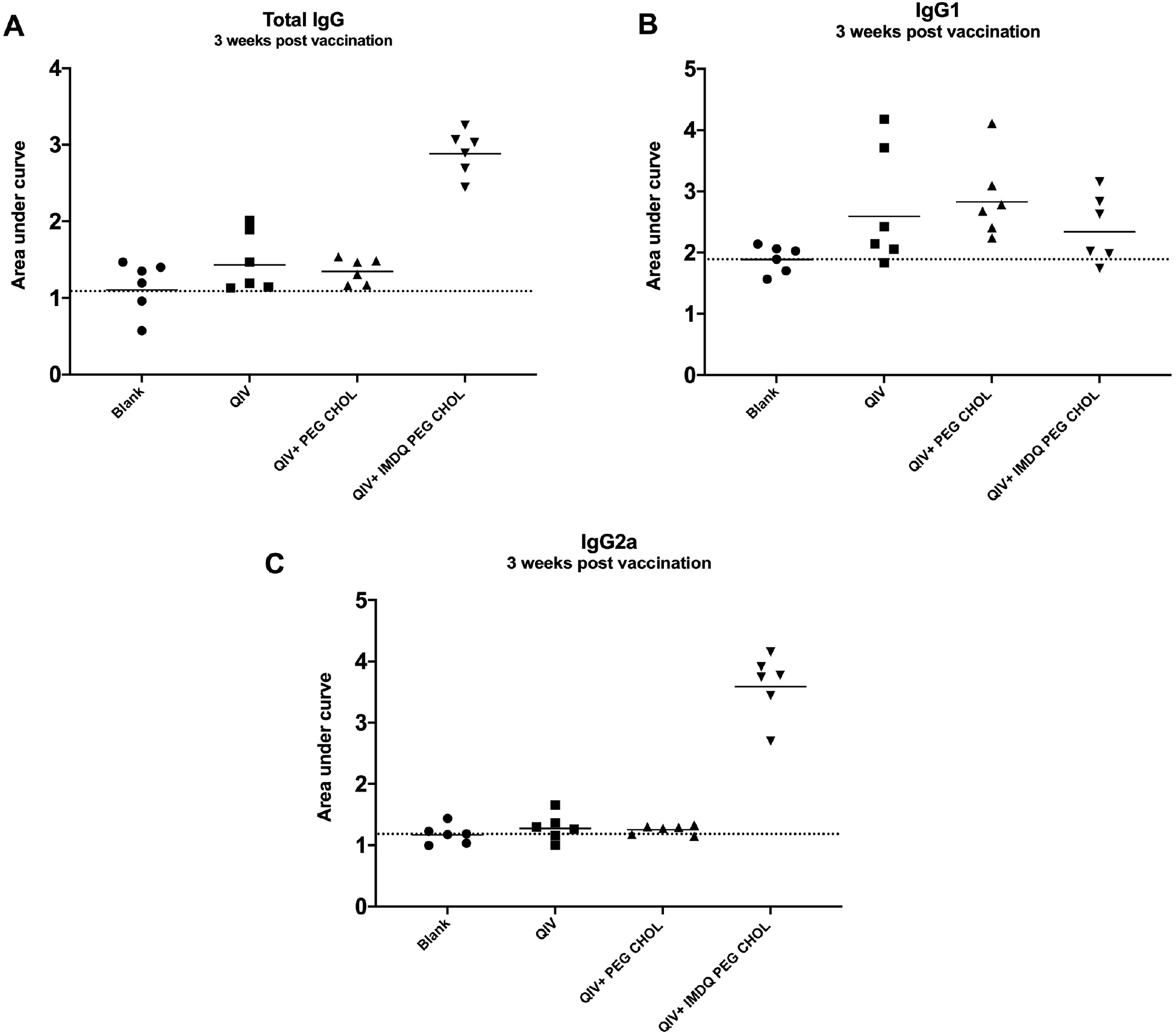
ELISA titers for total IgG, IgG1 and IgG2a in mice sera collected 3 weeks post-vaccination. The ratio IgG2a/IgG1 is representative of the Area under the OD450 curve of the individual serum sample.

**Supplementary Fig. 5:**
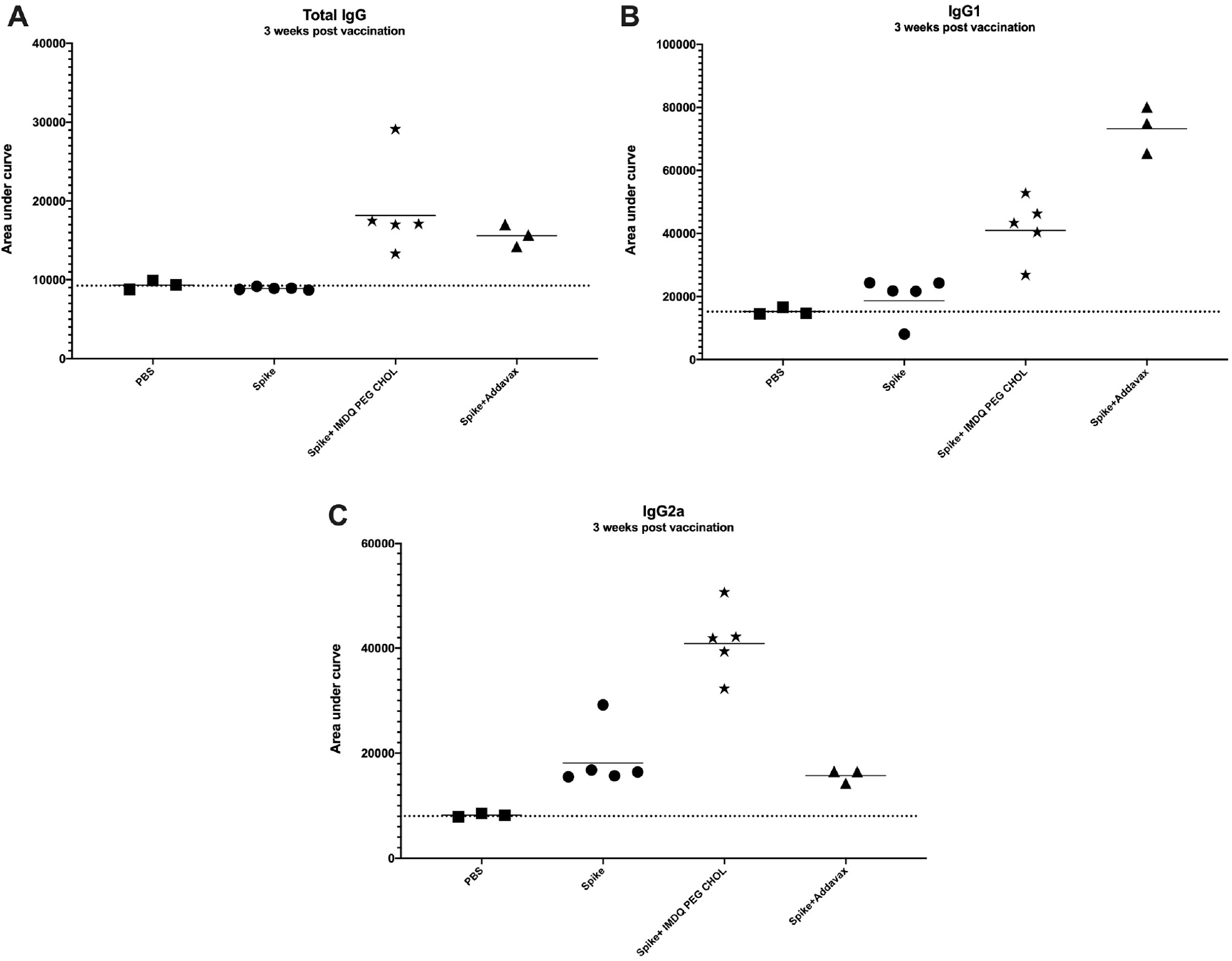
ELISA titers for total IgG, IgG1 and IgG2a in mice sera collected 3 weeks post-vaccination. The ratio IgG2a/IgG1 is representative of the Area under the OD450 curve of the individual serum sample.

